# Physical constrains and functional plasticity of cellulases: Linear scaling relationships for a heterogeneous enzyme reaction

**DOI:** 10.1101/2020.05.20.105569

**Authors:** Jeppe Kari, Gustavo A. Molina, Kay S. Schaller, Stefan J. Christensen, Corinna Schiano-di-Cola, Silke F. Badino, Trine H. Sørensen, Nanna Røjel, Malene B. Keller, Bartlomiej M. Kolaczkowski, Johan P. Olsen, Kristian B. R. M. Krogh, Kenneth Jensen, Ana Mafalda Cavaleiro, Günther H. J. Peters, Nikolaj Spodsberg, Kim Borch, Peter Westh

**Author notes:** Both authors contributed equally to this work (Jeppe Kari, Gustavo A. Molina).

## Abstract

Enzyme reactions, both in Nature and technical applications, commonly occur at the interface of immiscible phases. Nevertheless, stringent descriptions of interfacial enzyme catalysis remain sparse, and this is partly due to a shortage of coherent experimental data to guide and assess such work. We have produced and kinetically characterized 83 cellulases, which revealed a conspicuous linear free energy relationship (LFER) between the strength of substrate binding and the activation barrier. This common scaling occurred despite the investigated enzymes were structurally and mechanistically diverse. We suggest that the scaling reflects basic physical restrictions of the hydrolytic process and that evolutionary selection have condensed cellulase phenotypes near the line. One consequence of the LFER is that the activity of a cellulase can be estimated from substrate binding strength, irrespectively of structural and mechanistic details, and this appears promising for *in silico* selection and design within this industrially important group of enzymes. On a more general note, the LFER may identify a link to inorganic heterogeneous catalysis, and hence open for the implementation of approaches from this field within interfacial enzymology.

## Introduction

Enzyme reactions at interfaces are common both in Nature and industry ^1^. About half of the enzymes in the living cell work at a membrane surface ^2^ and many technical enzyme applications involve catalysis at the solid-liquid interface ^3^. Examples of the latter include the use of immobilized enzymes in protein arrays or biosensors ^4^, but more commonly, the activity of soluble enzymes on insoluble substrates such as polysaccharides, lipids or precipitated proteins ^5^. Studies of heterogeneous enzyme reactions have shown that both substrate specificity ^6^, turnover number ^7^, and enzyme-substrate binding affinity ^8^ can be significantly altered at an interface compared to analogous reactions in the bulk. Nevertheless, the kinetics of interfacial reactions is typically disregarded or fleetingly treated in textbooks ^9-13^, and this state of affairs is quite different from conventional (non-biochemical) catalysis, where homogeneous and heterogeneous reactions are treated in parallel. Although insightful models and concepts of interfacial enzyme kinetics have been suggested ^14-16^, no generally applied kinetic approach or rate equation currently exist. Neither is it clear whether progress in this field should be based on adaptation of conventional enzyme kinetic theory, or modifications of concepts and principles taken from inorganic heterogeneous catalysis.

In the current work, we investigate heterogeneous enzyme catalysis using cellulases as a paradigm. These enzymes catalyze the hydrolysis of the β-1,4 glycosidic bond that links glucopyranose units in (insoluble) cellulose and constitute a generic and experimentally convenient example of interfacial enzymes. In addition, cellulases are of direct industrial interest since enzymatic conversion of lignocellulosic biomass to fermentable sugars (known as saccharification) is expected to play a key role in the upcoming biorefineries that produce fuels, chemicals, and materials from sustainable feedstocks ^17-19^. We focused on fungal cellulases, which are commonly applied in industrial enzyme cocktails ^20^, and investigated enzymes from Glycoside Hydrolase (GH) family 5, 6, 7, 12 and 45 ^21^. Specifically, we produced and biochemically characterized 83 enzymes using insoluble cellulose as substrate. The characterized cellulases included both wild types and variants and represented a wide range of structural and functional differences (see Fig. 1). Nevertheless, the kinetic data showed a clear common trait as we found a conspicuous scaling between ligand binding strength and maximal turnover across the entire group of cellulases. The scaling could be expressed as a so-called linear free energy relationship (LFER), and we used this to discuss functional plasticity and physical constraints for the enzymatic conversion of cellulose. We argue that the LFER for cellulases may facilitate both mechanistic and evolutionary studies, and act as guidance in future attempts to select or design improved technical enzymes. Moreover, the LFER identified here for an interfacial enzyme reaction may establish links to the principles of inorganic heterogeneous catalysis, and hence help develop better theoretical frameworks for interfacial enzyme reactions.

**Fig. 1.**
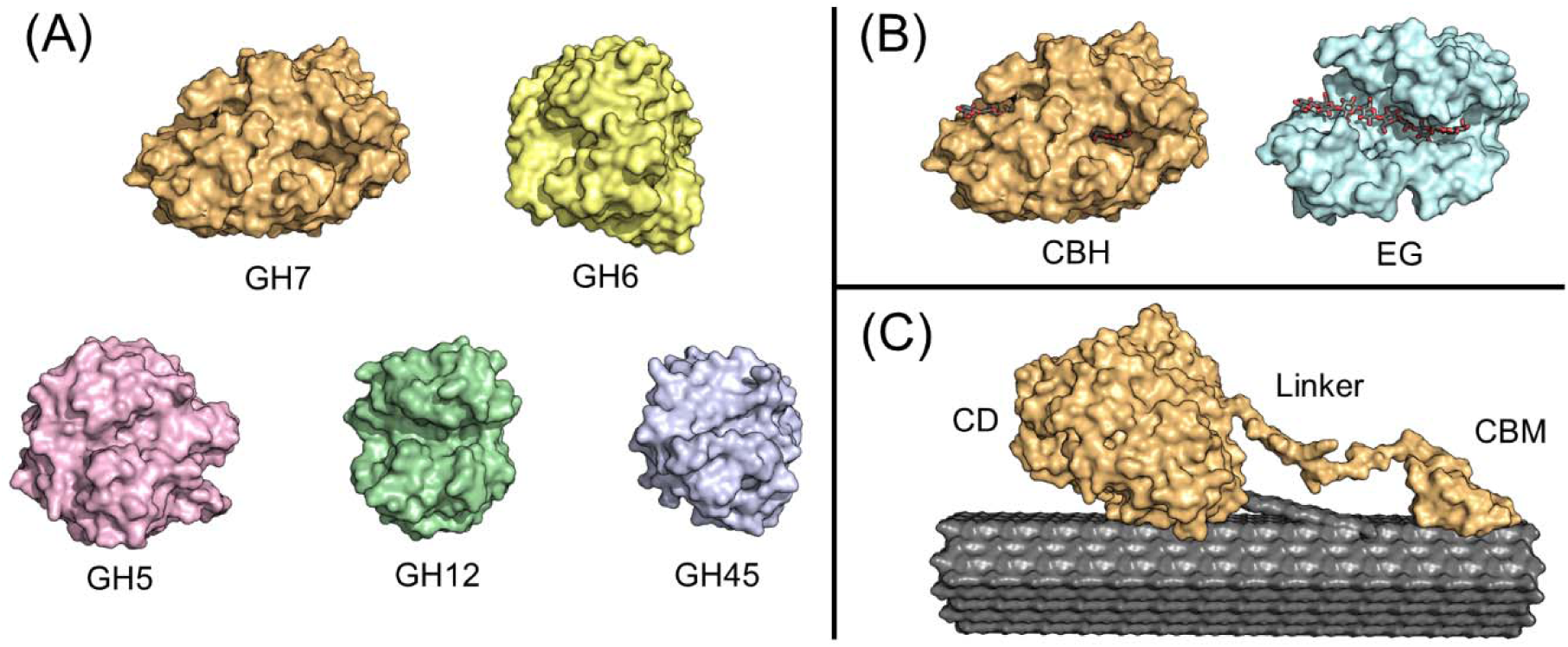
Structural representation of different classes of cellulases characterized in this study. A) Overall fold of the six different glycoside hydrolase (GH) families (PDB: 4C4C (GH7), 1QK2 (GH6), 1H8V (GH12), 4ENG (GH45), 3QR3 (GH5)). B) Structure of two GH7 cellulases with different modes of action in complex with cellononaose. An exo-active cellobiohydrolase (CBH) with a tunnel-shaped catalytic domain (PDB: 4C4C) and endo-active endoglucanase (EG) with an open catalytic cleft (PDB: 1EG1). C) Illustration of a GH7 CBH in complex with a cellulose fiber. The enzyme is modular with a catalytic domain (CD) and a carbohydrate-binding module (CBM) connected by a flexible linker. All structures were visualized using PyMOL (Schrödinger, L. (2015) The PyMol Molecular Graphics System. Version 2.3.0.).

## Results

### Enzyme production

The investigated enzymes were selected from five GH families as illustrated in Fig. 1 and Table 1. These families (GH5, GH6, GH7, GH12, and GH45) cover essentially all major fungal cellulases^20^ and hence represent a wide range of structures and mechanisms. This included enzymes with or without a carbohydrate-binding module (CBM), enzymes using an inverting or retaining mechanism, enzymes that attack the cellulose chain internally (endoglucanases, EGs) or at a chain end (cellobiohydrolases, CBHs) and enzymes with different degree of processivity. In addition to the wild type enzymes, a library of cellulase variants was made with the intention of changing the enzyme-substrate binding strength. This library included variants with mutations in the CBM, linker, and catalytic domain as well as variants where the CBM and linker were added, removed or swapped. A full list of the enzymes characterized here can be found in Supplementary Table 2.

**Table 1.** Fungal cellulases characterized in this study.

| GH family | Structural fold | Catalytic mechanism | Mode of action | EC number | Number of enzymes |  |
| --- | --- | --- | --- | --- | --- | --- |
|  |  |  |  |  | Wild-types | Variants |
| 7 | β-jelly roll | Retaining | CBH | 3.2.1.176 | 11 | 21 |
|  |  |  | EG | 3.2.1.4 | 3 | 6 |
| 6 | $\alpha/\beta$ barrel | Inverting | CBH | 3.2.1.91 | 5 | 16 |
| 5 | $(\beta/\alpha)_8$ | Retaining | EG | 3.2.1.4 | 8 | 0 |
| 12 | β-jelly roll | Retaining | EG | 3.2.1.4 | 3 | 4 |
| 45 | $\beta$ barrel | Inverting | EG | 3.2.1.4 | 6 | 0 |
| **Total** |  |  |  |  | **83** |  |

### Kinetic analysis

All enzymes were characterized by Michaelis-Menten (MM) kinetics using microcrystalline cellulose (Avicel PH-101) as substrate. Quasi-steady-state rates (*v*_*ss*_) were measured at a constant, low enzyme concentration (E_0_) and different substrate loads (S_0_), and analyzed by the MM-equation (Eq. 1) using non-linear regression. The resulting kinetic parameters (K_M_ and *k*_*cat*_) are listed in Supplementary Table 2.

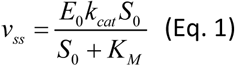

Previous studies have identified practical procedures for measuring the quasi-steady-state rate for this type of system ^23^ and shown that Eq. 1 is valid and applicable even though the substrate is solid and specified by its mass load (S_0_) in units of g/L ^24,25^. The derived rates were based on soluble products only and control experiments (Supplementary Table 1) showed that this was a good descriptor of the overall activity even for EGs.

We used *k*_*cat*_ and K_M_ to estimate changes in respectively transition-state free energy (ΔΔ*G*^‡^) and standard free energy of ligand binding 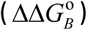 (c.f. Fig. 5) following well-established principles ^26,27^. Specifically, we used the equations

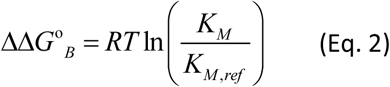

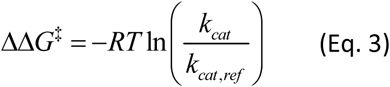

Equations 2 and 3 introduce a reference enzyme with the kinetic parameters K_M,ref_ and *k*_*cat,ref*_, and hence the calculated free energies are energy changes relative to the selected reference. This approach alleviates ambiguities regarding standard states (Eq. 2) and pre-exponential factors (Eq. 3). We used the GH6 cellobiohydrolase from *Trichoderma reseei* (TrCel6A) as our reference enzyme and it follows that this enzyme will have ΔΔ*G*^‡^ = ΔΔ*G*° _*B*_= 0 (*i.e.* TrCel6A will be located in the origin of Fig. 2 below). We emphasize that the selection of reference enzyme merely serves to define an origin; it has no bearing on the subsequent discussion, which considers changes in the free energy.

**Fig. 2.**
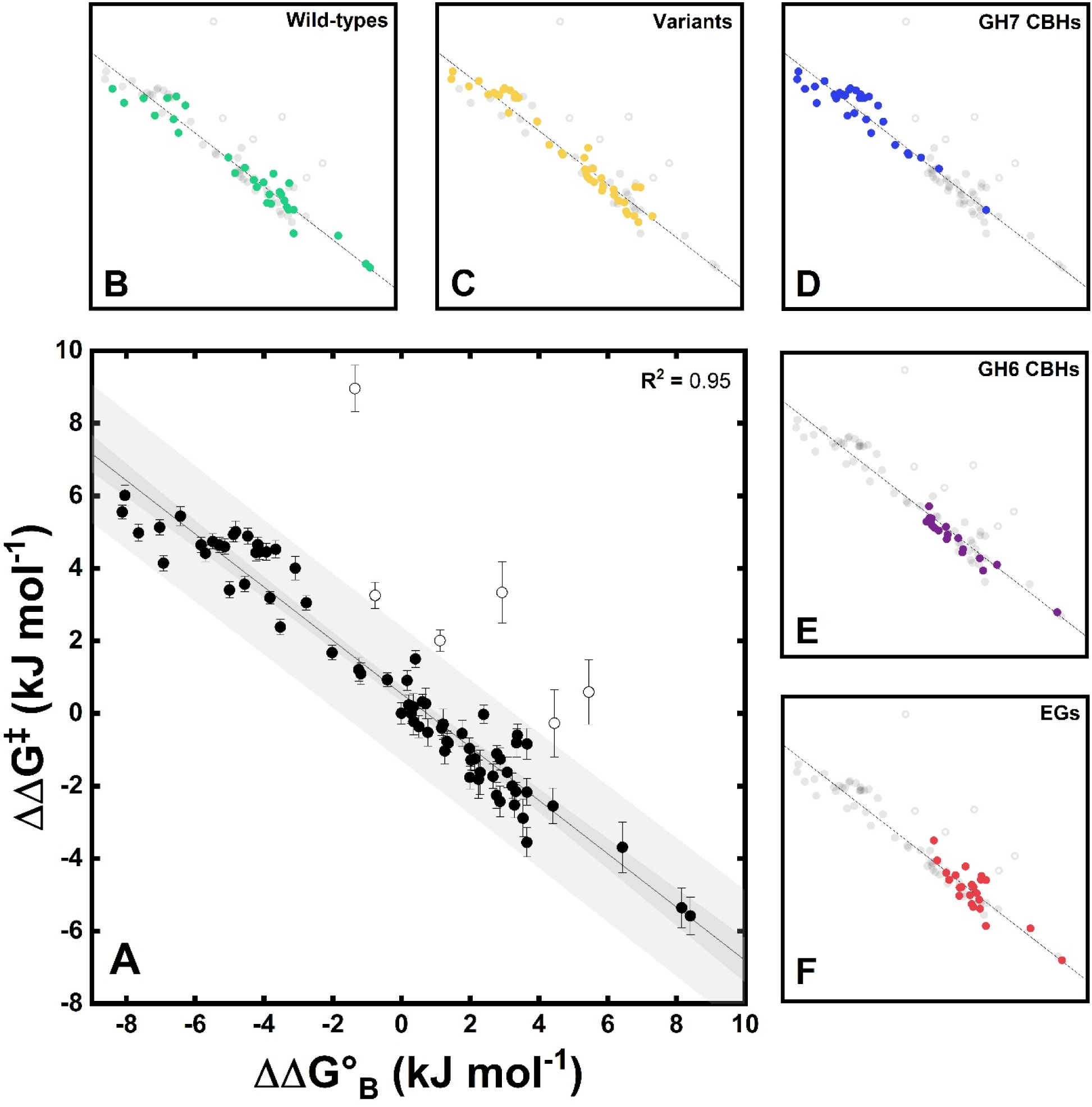
Correlation plot of change in the binding free energy (ΔΔG°_B_, Eq. 2) and activation free energy (ΔΔG ^‡^, Eq. 3) for all investigated enzymes (A). The smaller panels highlight data for different classes of enzymes. These are wild type cellulases (B), variants (C), cellobiohydrolases from GH7 (D), cellobiohydrolases from GH6 (E), and endoglucanases from family GH7, GH12, and GH45 (F). The solid line in all plots derives from the linear regression to all the experimental data of the main panel (A) excluding the outliers (open symbols) identified as explained in the main text. Bands shown in panel A are 95% confidence band (dark gray) and 95% prediction band (light gray).

In Fig. 2, the change in activation free energies (ΔΔ*G* ^‡^) are plotted against the change in standard binding free energies (ΔΔ*G*°_*B*_) for all investigated enzymes. From the main panel (Fig. 2A), it appeared that most enzymes clustered in a narrow lane around the diagonal. Some enzymes were located above the diagonal, but we did not find any below. To assess whether the experimental points in Fig. 2 correlated with structural or functional properties of the studied enzymes, we made five other plots (Panels B-F, Fig. 2) that highlighted specific groups in the dataset. Linear regression showed that the slope in Fig. 2 was – 0.74±0.02. Regression outliers were identified based on studentized residual analysis using a conservative cutoff of ± 2.5σ. The outliers were omitted from the regression analysis and identified by open symbols in Fig. 2. A list of kinetic parameters for all investigated enzymes can be found in Supplementary Table 2.

### Computational analysis

The strong correlation between binding- and activation- free energies shown in Fig. 2, may open up for prediction of catalytic rates based solely on computed binding free energies. To test this approach, we calculated cellulose binding strengths for a subset of nine enzymes from Fig. 2, using molecular dynamics (MD) simulations with umbrella sampling along the reaction path. We selected enzymes that spanned a wide range of ΔΔG°_B_ and represented all structural and functional classes listed in Fig. 1. For modular cellulases, the contribution of the CBM to ΔΔG°_B_ was computed separately. Comparisons in Fig. 3 showed that despite the diversity of the analyzed cellulases, computed changes in binding affinity, ΔΔ*G*_*B,MD*_, scaled reasonably with the experimental values, ΔΔ*G*°_*B,Exp*_.

**Fig. 3.**
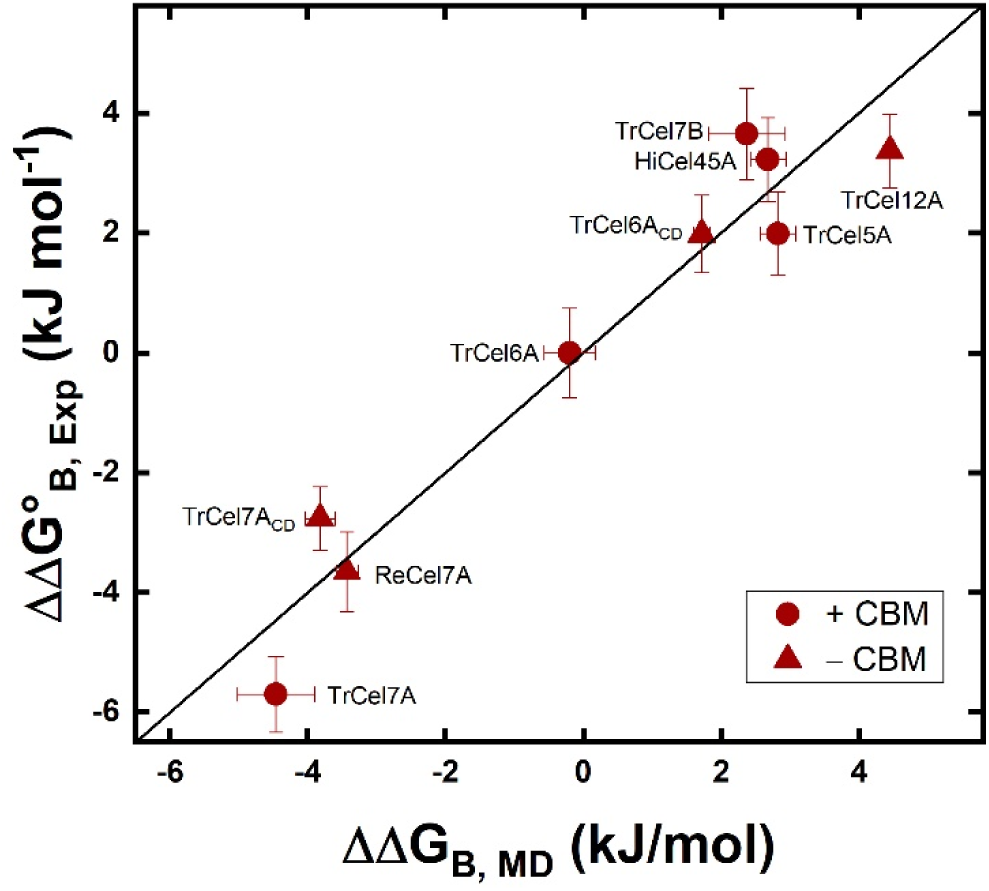
Correlation of computed (ΔΔG_B,MD_) and experimental (ΔΔG° _B_) changes in free energies of binding for nine selected cellulases. Experimental values are taken from Fig. 2. The selected cellulases covered a wide range of ΔΔG°_B_ shown in Fig. 2 and encompassed all main structural and functional traits specified in Fig. 1.

## Discussion

We have produced and kinetically characterized 83 enzymes covering essentially all classes of fungal cellulases (Table 1 and Fig 1). We used the same expression host, to ensure the enzymes were exposed to the same apparatus of post-translational modifications. Moreover, kinetic characterizations were based on the same substrate, experimental conditions, and principles of analysis. This provided an unprecedented basis for comparative analysis of interfacial enzymes in general and cellulases in particular. Indeed, the breadth of the data-set allowed us to identify a striking LFER between ΔΔ*G*^‡^ and ΔΔ*G*°_*B*_, and in the following we discuss origins and corollaries of this observation.

### Enzyme fitness and physical constraints

Fig. 2 presents a fitness landscape for cellulases attacking their native insoluble substrate, and it appears that most enzymes accumulated around the diagonal. This defines a continuum of properties ranging from enzymes with high substrate affinity but a low turnover rate (upper left end, Fig. 2A), to enzymes with a low substrate affinity but a high turnover number (lower right end, Fig. 2A). The tendency to accumulate along the diagonal was observed for all types of cellulases (refer to Table 1 and Fig. 1), and hence does not seem to rely on specific structural or mechanistic properties. Rather, it appears that the maximal turnover can be expressed solely by one descriptor, namely substrate affinity. The area above the diagonal in Fig. 2 represents a combination of low turnover and weak ligand binding, and this seems to signify inefficient catalysis. We found some enzymes in this range, including some wild type enzymes and variants with replacements of key amino acid residues. We suggest that this northeastern region of the fitness landscape represents enzymes that have been either catalytically impaired by our engineering, structurally unstable, or have other primary substrate preference than cellulose.

On the other hand, the region below the diagonal in Fig 2, specifies enzymes which have both strong substrate binding and rapid turnover (both ΔΔ*G*°_*B*_ and ΔΔ*G*^‡^ are low compared to the reference). These traits clearly appear functionally advantageous, but we did not find any cellulases in this region. We suggest that this absence is the result of basic physical restrictions of the cellulolytic process. It follows that the accumulation of data points in a narrow lane in Fig. 2 may be seen as a balance between evolutionary selection, which drives the kinetic parameters towards the southwest, and physical constraints, which prevents this development beyond the boundary defined by the line in Fig. 2.

The engineered variants in Fig. 2B represents a range of replacements and deletions at different positions (see Supplementary Table 1), which were designed with the overall purpose of altering ligand binding strength. In a few cases, the mutations shifted the variants into the northeastern wasteland of the fitness landscape, but most remained on the diagonal. The tendency to stay on the line did not reflect that the variants had similar kinetic parameters. Rather, changes in K_M_ and *k*_*cat*_ tended to compensate. Some examples of this are highlighted in Fig. 4, and it appears that both point mutations and extensive changes in the amino acid sequence readily moved kinetic parameters up or down the diagonal, but rarely send them off the line. Interestingly, the vast majority of the variants moved down the line compared to their respective wild type, and only in cases where a CBM was added to a CBM-less wild type (Fig. 4A) did the variant move up the line towards higher affinity and lower turnover. This indicates that cellulases have evolved a fine-tuned substrate affinity to cover a specific affinity window. This quality may be important in nature where cellulose is degraded by multiple types of cellulases with different affinities.

**Fig. 4.**
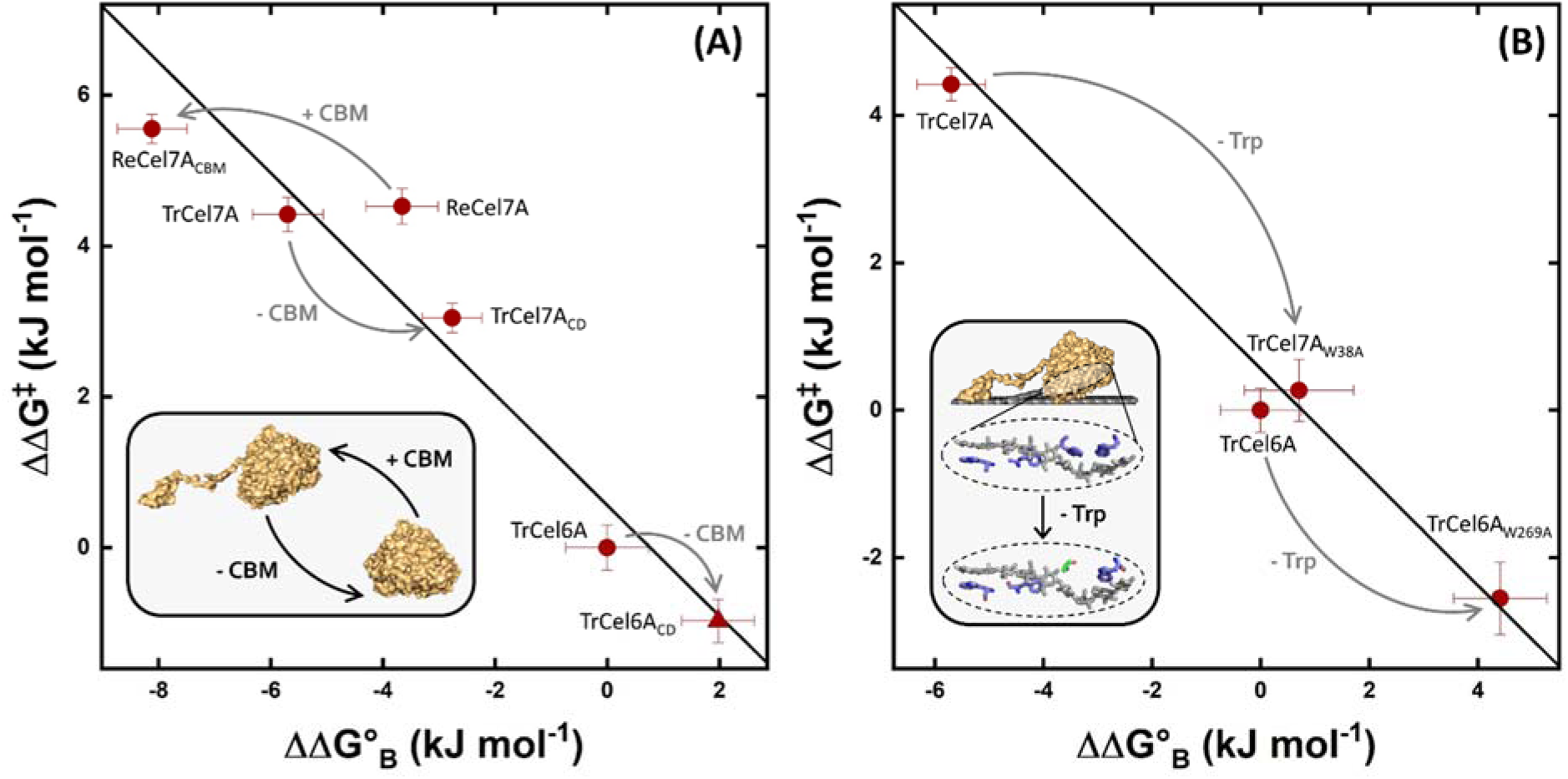
Illustration of the effect of the non-catalytic CBM (A) and tryptophan residues in the catalytic domain of cellobiohydrolases from GH6 and GH7 (B). Correlation plot of ΔΔG° _B_ and ΔΔG^‡^ for 3 wild type CBHs and 3 variants, where the CBM was either removed (-CBM) from the wild type (TrCel7A → TrCel7A_CD_, TrCel6A → TrCel6A_CD_) or added (+CBM) to the wild type (ReCel7A → ReCel7A_CBM_). Analogous correlation plot for replacements of conserved tryptophan residues by alanine in the catalytic domain of TrCel7A (TrCel7A → TrCel7A_W38A_) or TrCel6A (TrCel6A → TrCel6A_W269A_). The solid line shown in both plots is the same as in Fig. 2. It appears that changes in ΔΔG° _B_ and ΔΔG^‡^ tend to compensate so that all enzymes remain close to the diagonal. Inserts are illustrations to guide the reader about the structural changes in the variants.

### Origin of physical constrain

Correlations between binding and activation free energies are well-known in both organic and inorganic catalysis ^28^ but have only been sporadically used for (homogenous) enzyme reactions ^29,30^. A linear free energy relationship exists if the binding free energy, ΔG^°^_B_, scales linearly with the free energy of activation, ΔG^‡^. This is tantamount to proportionality between the changes in these two functions, and we may write

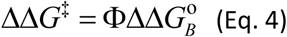

The proportionality constant, Φ (slope in Fig. 2) provides some information about the nature of the transition state (TS), and this has been used, for example, to elucidate the TS of protein folding ^31^. As proposed by Warshel ^27^, the Φ-value also provides a means to classify effects of mutations on enzyme function. If, for example, both the enzyme-substrate complex and the transition state in a variant are stabilized to the same extent (so-called uniform binding, see Fig. 5 B-1) Φ would be 0 since the activation energy would remain unchanged (i.e. ΔΔ*G*^‡^ = 0). Another illustrative case is when changes in interactions only manifest themselves in the TS (so-called TS stabilization, Fig. 5B-2). This results in Φ →∞ since the activation energy can be changed independently of the binding energy. Finally, if mutations only act to stabilize the ground state complex (GS stabilization, Fig. 5B-3), ΔΔ*G*^‡^ will change commensurate with ΔΔ*G* ° *B*, and Φ = −1.

This interpretation of Φ-values was developed to classify mutants, but in the current context it may also elucidate differences across the investigated group of cellulases (wild types and mutants). From Fig. 2 we found Φ = −0.74 ± 0.02, and it follows that kinetic differences among the investigated cellulases can be mostly ascribed to differences in the degree of GS stabilization. This has the noticeable consequence that the free energy of the (rate-limiting) TS is quite similar for all tested enzymes, and that the main kinetic diversity lies in different affinities for the substrate. Experimental studies have suggested that the rate-limiting step for some cellulases is slow dissociation ^32-36^. Since weaker binding is associated with a lower activation barrier for dissociation (Fig 5B-3), a dissociation limited mechanism would explain the inverse correlation of binding strength and maximal turnover. Based on these considerations it is tempting to suggest that weak ligand binding is a functional advantage since *k*_*cat*_ increases. However, mutational studies suggest that this is not the case for glycoside hydrolases attacking solid carbohydrates ^37,38^. The characterized variants support this interpretation since most of the variants moved down the line in Fig. 2 compared to the respective wild type, indicating that the wild type was optimized for high affinity. Strong ligand binding may be needed in order for the enzyme to transfer a cellulose chain from the cellulose surface, where it is strongly bound ^39,40^, to the binding cleft (see cartoon in Fig. 5A). Hence, strong ligand binding appears to benefit catalysis by promoting ligand transfer ^41^, but it is inevitably associated with a slow turnover of an off-rate controlled reaction, as illustrated in Fig. 5 B-3.

### Consequences of observed LFER

One aspect of the proposed scaling of K_M_ and *k*_*cat*_ is that the initial rate, *v*_ss_ (Eq. 1), may be described by just one of the kinetic parameters. To illustrate this, we combined eqs. 2-4 and solved for *k*_*cat*_.

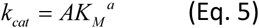

In eq. (5) *a* = −Φ and 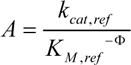. Inserting eq. 5 into the MM-equation (Eq. 1) expresses *v*_ss_ as a function of K_M_

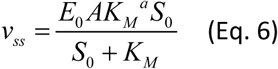

Eq. 6 underscores how ligand affinity is a double-edged sword. Hence, as demonstrated in the SI, eq. 6 has a global maximum when K_M_ attains the value

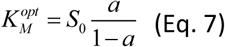

This implies that at a fixed load of substrate, S_0_, a cellulase with low K_M_ (*i.e.* 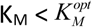) will become a better catalyst (increase *v*_*ss*_) if it is engineered for weaker substrate binding. Conversely, a weakly binding enzyme 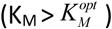 will gain from tighter binding. In the current case, *a* = 0.74 and insertion into eq. (7) shows that 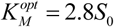. In other words, the fastest initial rate on the current substrate (Avicel) will be a cellulase that has a K_M_ value which is around three-fold higher than the Avicel load. To illustrate this, we plotted *v*_*ss*_ as a function of K_M_ for all the investigated enzymes (excluding outliers identified in Fig. 2) at different substrate loads (Fig. 6). The results are in line with a previous observation ^42^ showing the so-called volcano plots, where cellulase activity tapers off on each side of the optimal affinity. Such volcano plots mirror the Sabatier principle, which states that the catalytic efficacy is optimal for a catalyst with intermediate binding strength ^43^. Higher/lower affinity leads to a situation where dissociation/association limits the overall rate. The optimal affinity, 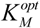, depends on the substrate load and this is indicated by the black symbols in Fig. 6, which were calculated using eq. 7. We emphasize that the appearance of an optimal K_M_ is a direct consequence of the LFER in Fig. 2, and that this type of analysis is well established within (non-biochemical) heterogeneous catalysis ^44,45^.

**Fig. 5.**
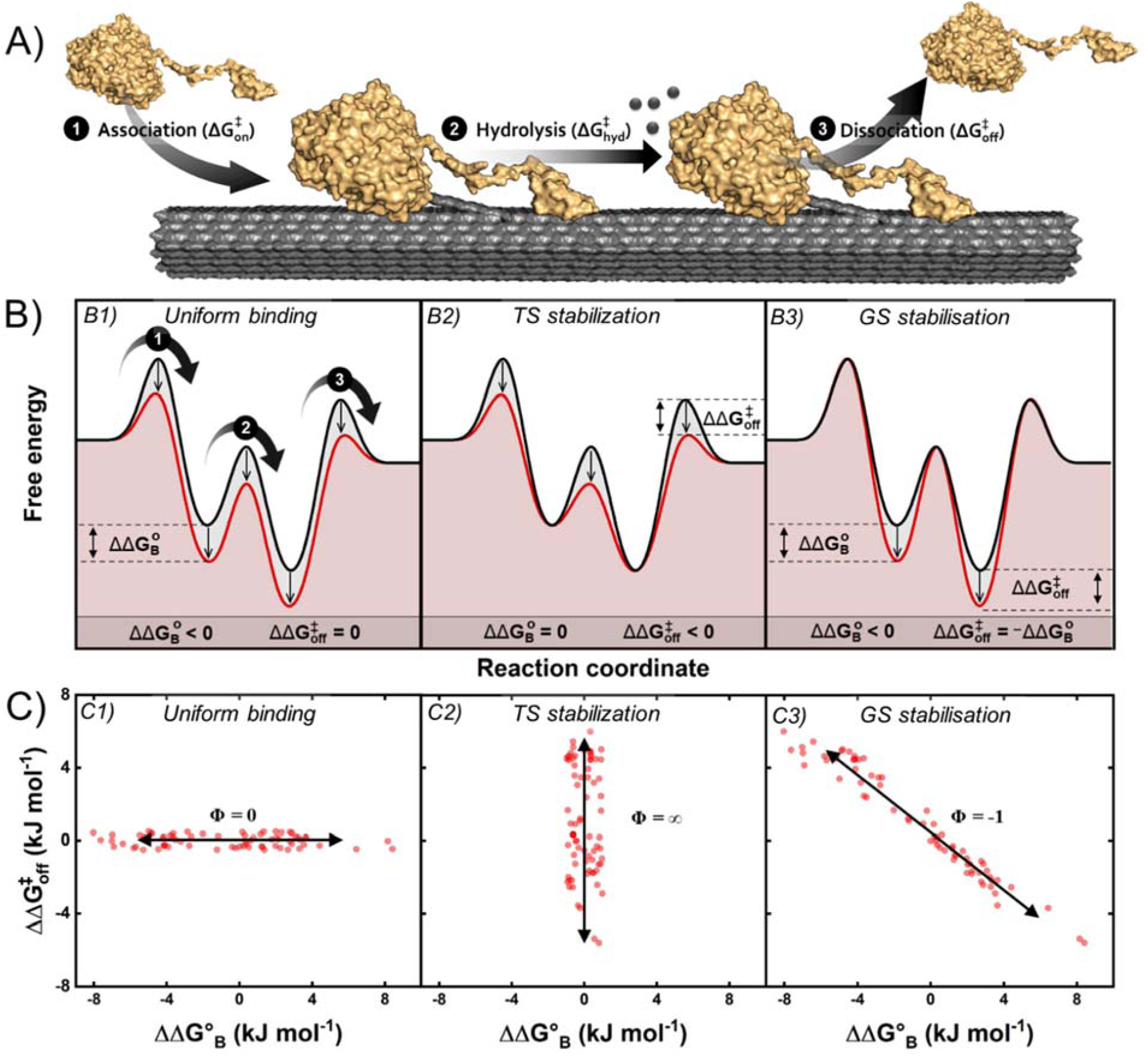
Origin of the physical constrain on cellulose. (A) Simplified reaction scheme for a cellulase (yellow) hydrolyzing insoluble cellulose (gray). The cartoon provides a structural interpretation of three steps in the overall reaction; (1) association, (2) hydrolysis, and (3) dissociation. (B) Schematic energy diagrams for a wild type (black curve) and three conceptually different variants (red curves). (C) Expected scaling plots for a group of variants that behave according to the three different energy-diagrams shown in panel B. If the energy of the variant differs from the wild type by the same amount in both transition state (TS) and ground state (GS), we have so-called uniform binding and Φ= 0 (panel B1 and C1). The parallel shift in energies for uniform binding implies that the same interactions occur in GS and TS. If, on the other hand, mutation only lowers the TS energy, it is denoted TS-stabilization, and this corresponds to a vertical line in the scaling plot (panel B2 and C2). Finally, in GS-stabilization (panels B3 and C3), only the GS energy changes, while the TS remains fixed. In this case, Φ= −1, and this is close to the experimental behavior observed in Fig. 2.

**Fig. 6.**
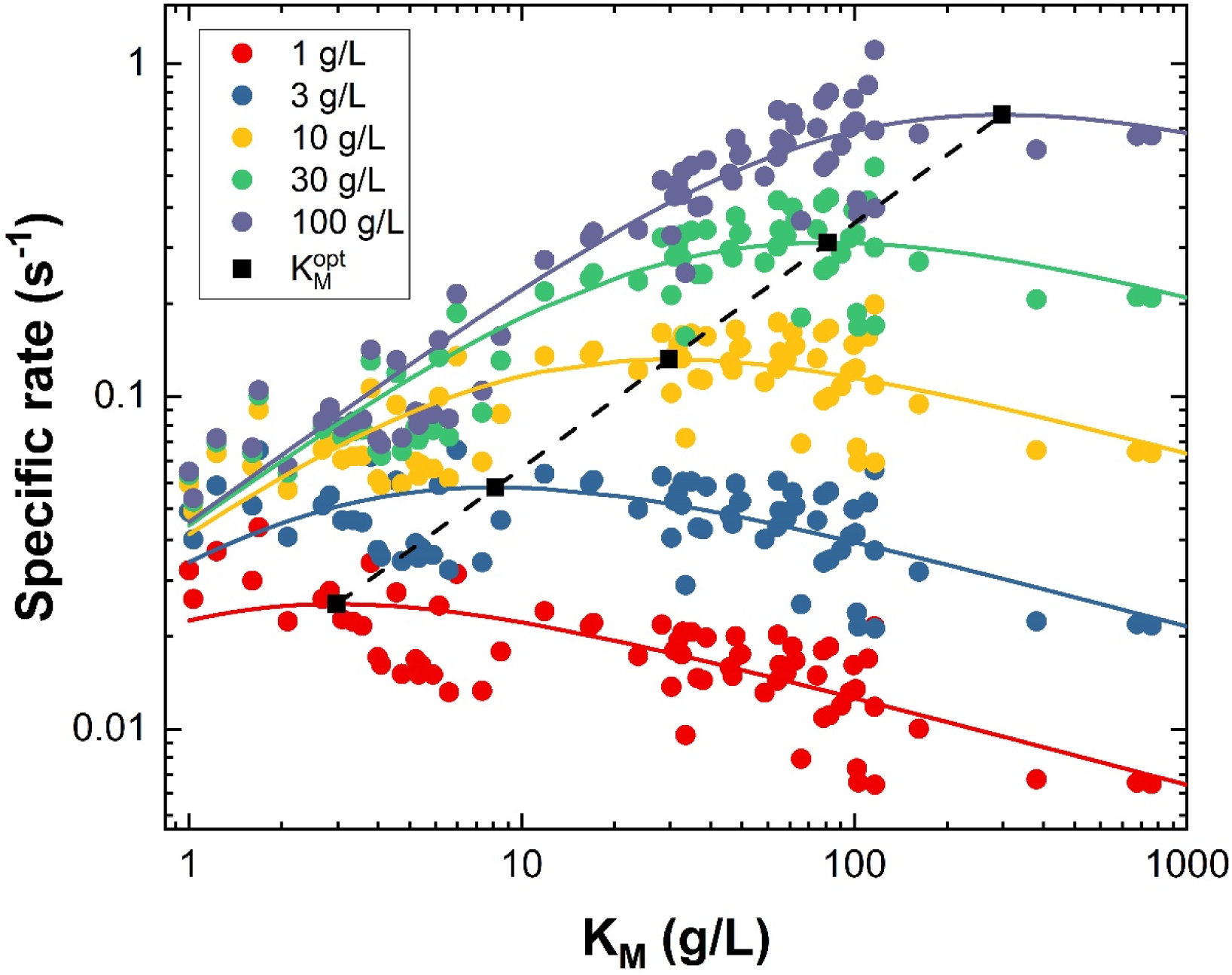
Volcano Plots showing the specific rate as a function of K_M_ for the investigated enzymes (excluding outliers identified in Fig. 2) at five different substrate loads. Points represent experimental data and solid lines are the predicted volcano curves calculated using Eq. 6 and a = 0.74 (there are no free parameters in the determination of these curves). Black squares represent 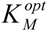 values calculated using eq. (7), and these points identify the maxima of the volcano plots at a given load of substrate.

As a final example of an application of the LFER in Fig. 2, we note that it may be useful in computational selection and design of enzymes for technical use. Thus, a link between activity and affinity provides an important simplification as it converts the highly complex problem of *in silico* assessment of enzyme turnover frequency to the more tractable challenge of calculating binding energy. To illustrate this, we computationally assessed the strength of enzyme-substrate interactions for a subset of nine enzymes spread along the diagonal in Fig. 2. As shown in Fig. 3, the computed binding energies scaled with the experimental values. These results suggest that the kinetic properties of novel, uncharacterized enzymes may be estimated by combining computed binding data with an experimental LFER based on a limited number of enzymes. Hence, efficient enzymes for a given set of experimental conditions could be identified through *in silico* screening.

### Concluding remarks

Kinetic characterization of a wide group of fungal cellulases on their native, insoluble substrate revealed a linear free energy relationship between substrate binding and activation barrier. We propose that this reflects basic physical restrictions of the hydrolytic reaction, which limits the evolutionary selection to a narrow lane around the scaling line, irrespectively of the enzymes’ fold, modularity, or catalytic mechanism. The scatter around the proposed scaling line was about 2 kJ/mol, and in the logarithmic scale of Fig. 2, this corresponds to a factor of about 2 in the value of *k*_*cat*_ (see eq. 3). Hence, our results suggested that experimental *k*_*cat*_ values for enzymes with approximately the same K_M_ varied within this range. This variance encompassed a minor contribution from experimental errors (Supplementary Table 1), but it may also reflect kinetic diversity that results from differences in the mechanism and specificity of the tested enzymes. However, when we zoomed out and considered a broad range of K_M_-values, this variance was modest, and the fitness landscape was dominated by a common scaling for all enzymes. Comparisons of wild types and variants revealed that small alterations in sequence could lead to significant kinetic changes. In most cases, however, the changes involved a stringent movement up or (more commonly) down the scaling line rather than a shift away from the line, and this further demonstrated a strong coupling between affinity and turnover. We propose that this behavior is linked to the interfacial nature of the reaction. On one hand, strong ligand interactions are required to enable the transfer of a cellulose chain from the cellulose surface to the enzyme complex. On the other, a highly stable enzyme-substrate complex is inescapably associated with slow turnover (Fig. 5 B-3). These relationships may help rationalize cellulolytic mechanisms and guide the selection of technical enzymes. It also appears that LFERs for interfacial enzyme reactions may establish a new connection to (inorganic) heterogeneous catalysis, and hence pave the way for the use of practices and principles from this field within enzymology.

## Methods

### Enzymes and kinetic measurements

Experimental methods used in this work have been described elsewhere (see Supplementary Table 2). Briefly, we expressed all enzymes heterologously in *Aspergillus oryzae* and purified as described elsewhere ^46,47^. SDS-PAGE gels (15-well NuPAGE 4–12% BisTris, GE Healthcare) revealed a single band for the purified enzymes, and their concentrations were determined by UV absorbance at 280 nm using a theoretical extinction coefficient calculated based on amino acid sequence 48. Michaelis-Menten curves were obtained as described previously ^47^ using 0.1 µM enzyme and microcrystalline cellulose (Avicel PH-101, Sigma-Aldrich, St. Louis, MO) load ranging from 1 to 100 g/L. All experiments were done at 25 °C using standard buffer 50 mM sodium acetate pH 5.0 in triplicates.

### Molecular dynamics simulation

For simplicity, the modular structures of the different cellulases were split into two simulations. The CDs were simulated in complex with a cellononaose ligand and the CBMs (if present) were simulated bound to a cellulose crystal.

### Simulations of the Catalytic Domains

If available, the structures were taken from the Protein Data Bank (TrCel7A: 4C4C, TrCel7B: 1EG1, TrCel6A: 1QK2, TrCel12A: 1H8V, TrCel45A: 3QR3, HiCel45A: 4ENG, Re7A: 3PL3). The ligand was inserted by alignment, if a related structure with a similar large disaccharide was available. Elsewise, docking with Autodock Vina was performed ^49^. The ten clusters lowest in energy were inspected and the lowest energy configuration from the cluster with the closest distance between the catalytic residues and the glycosidic bond of interest was taken. The CHARMM36 force field was used to describe the system ^50^. All simulations were run in GROMACS 2018.6 ^51^. Catalytic acids of all CDs were protonated. GROMACS was used to construct a cuboid box with edge lengths of 9.4×9.4×20 nm and the complexes were positioned at 4.7, 4.7, and 3.3 nm. The complexes were rotated so that the center of mass of the last and the fourth last sugar unit of the ligands were parallel to the z-axis. The systems were solvated with TIP3P water. To neutralize the net charges of the systems, random water molecules were exchanged with sodium ions. Minimization was conducted in a steepest-descent over 10’000 iterations. All subsequent simulations were performed at 300K. *NVT*-simulations were performed for 100 ps while keeping the complex restraint. Thereafter, *NPT*-simulations with restraints on the solutes were performed for 100 ps. For all further simulation, only *C*_*α*_ further away than 1.5 nm from the ligand were restrained. A second round of *NPT*-simulations with the new restrains were performed for 100 ps. RMSD analysis of the protein backbone showed, that this time was sufficient to reach an equilibrated state. Thereafter, steered MD simulations were done over 800 ps with a pulling rate of 0.01 nm/ps and a force constant of 1000 kJ/mol/nm^2^. The pull was performed on the first sugar unit of the cellononaose ligand in z direction. The resulting trajectories were used to prepare further simulations. Frames every travelled 0.5 Å by the ligand were extracted up to a final distance of 1 nm between the CD and the ligand. The extracted frames were used as starting configuration for Umbrella sampling simulation along the binding path. Each window was simulated for 620 ps, where the first 20 ps were disregarded as equilibration. It should be noted, that TrCel6A works from the opposite end compared to the other cellulases ^20^.The set-up was adapted accordingly.

### Simulations of the Carbohydrate-Binding Modules

If available, the structures were taken from the Protein Data Bank (CBM1 of TrCel7A: 2CBH, CBM1 of TrCel7B: 4BMF). Otherwise, they were prepared through homology modelling by Modeller ^52^ (CBM1 of TrCel6A, CBM1 of TrCel5A, CBM1 of HiCel45A). A cellulose crystal of the type Iβ with a length of 5, a width of 6, and a depth of 3 unit cells was generated with the Cellulose Builder web server ^53^. The CBMs were placed on the surface according to Beckham, et al. ^54^. A cubic box with a minimal distance of 1.0 nm was constructed. The crystal plane was oriented perpendicular to the z-axis. The simulations were performed in a similar fashion than the ones for the CD domains. However, the heavy atoms of the crystals were kept constrained after the energy minimization and the second NPT-simulation was increased to 1 ns to get the CBM settled on the crystal surface.

### Analysis

Analysis of the trajectories was performed with GROMACS. The weighted histogram analysis method (WHAM) was applied to analysis the Umbrella sampling simulations along the binding path ^55^. If density gaps occurred, additional windows at those distances were inserted iteratively until no gaps occurred. From the resulting PFM curves, the energy difference between the minimum and the maximum of those curves were taken. The errors were estimated with bootstrapping. A linear regression of the experimental binding energy and experimental activation energy against the computer binding energy were performed. The former resulted in a linear fit in the form of y=0.16x+0.2 and with a Pearson’s coefficient r^2^=0.93 and the later resulted in y=0.13x+0.67 with r^2^=0.81. To counteract known overestimation issues of the method ^56-58^, a linear transformation on the initially obtained computed binding energies was performed using the parameters from the linear regressions. The final results for the prediction of the binding energies had a root-mean-squared error (RMSE) of 0.86 kJ/mol, the ones for the prediction of the activation energy had RMSE of 1.20 kJ/mol.

## Supporting information

sup. mat.

## Author contributions

JK and PW planned and designed the study. GAM characterized the enzymes, analyzed the data and interpreted the results along with JK. THS, AMC, NS, NR and KJ carried out cloning and expression of the enzymes. SJC, CSdC, SFB, NR, MBK, BK, JPO and KBRMK purified the enzymes, performed initial biochemical experiments, designed variants and selected wild type enzymes (also JK and GAM). KB aided in the selection of enzymes, development of hypotheses and data analysis. GHJP and KSS performed the MD simulations. JK, GAM, KSS and PW wrote the manuscript. All authors reviewed and approved the manuscript.

## Acknowledgements

This work was supported by Innovation Fund Denmark [Grant number: 5150-00020B], the Novo Nordisk Foundation [Grant number: NNF15OC0016606 and NNFSA170028392] and Independent Research Fund Denmark [Grant number: 8022-00165B].

## References

1 McLaren, A. D. & Packer, L. Some aspects of enzyme reactions in heterogenous systems. Adv. Enzymol. 33 (1970).

2 Berg, O. G. & Jain, M. K. Interfacial enzyme kinetic. (John Wiley & Sons, Ltd, 2002).

3 Basso, A. & Serban, S. Industrial applications of immobilized enzymes—A review. Molecular Catalysis 479, 110607, DOI:https://doi.org/10.1016/j.mcat.2019.110607 (2019).

4 Laurent, N., Haddoub, R. & Flitsch, S. L. Enzyme catalysis on solid surfaces. Trends Biotechnol. 26, 328–337, DOI:10.1016/j.tibtech.2008.03.003 (2008).

5 Kirk, O., Borchert, T. V. & Fuglsang, C. C. Industrial enzyme applications. Current Opinion in Biotechnology 13, 345–351, DOI:https://doi.org/10.1016/S0958-1669(02)00328-2 (2002).

6 Horan, N., Yan, L., Isobe, H., Whitesides, G. M. & Kahne, D. Nonstatistical binding of a protein to clustered carbohydrates. 96, 11782–11786, DOI:10.1073/pnas.96.21.11782 % J Proceedings of the National Academy of Sciences (1999).

7 Brode, P. F. & Rauch, D. S. Subtilisin BPN’: activity on an immobilized substrate. Langmuir 8, 1325–1329, DOI:10.1021/la00041a014 (1992).

8 Schurr, J. M. & McLaren, A. D. Enzyme action: comparison on soluble and insoluble substrate. Science 152, 1064–1066, DOI:10.1126/science.152.3725.1064 (1966).

9 Purich, D. L. Enzyme Kinetics: Catalysis & Control. 920 (Elsevier, 2010).

10 Cook, P. F. & Cleland, W. W. Enzyme kinetics and mechanism. (Garland Science, London, 2007).

11 Cornish-Bowden, A. Fundamentals of enzyme kinetics, 4th Edition. (John Wiley and Sons, 2012).

12 Segel, I. H. Enzyme kinetics: Behavior and analysis of rapid equilibrium and steady-state enzyme systems. Vol. 957 (Wiley, New York, 1975).

13 Fersht, A. Enzyme structure and mechanism. 2 edn, (W.H. Freeman, 1985).

14 Kartal, O. & Ebenhoh, O. A generic rate law for surface-active enzymes. FEBS Lett 587, 2882–2890, DOI:10.1016/j.febslet.2013.07.026 (2013).

15 Deems, R. A. Interfacial Enzyme Kinetics at the Phospholipid/ Water Interface: Practical Considerations. Analytical Biochemistry 287, 1–16, DOI:https://doi.org/10.1006/abio.2000.4766 (2000).

16 Gutiérrez, O. A., Chavez, M. & Lissi, E. A Theoretical Approach to Some Analytical Properties of Heterogeneous Enzymatic Assays. Analytical Chemistry 76, 2664–2668, DOI:10.1021/ac049885d (2004).

17 Chandel, A. K., Chandrasekhar, G., Silva, M. B. & Silvério da Silva, S. The realm of cellulases in biorefinery development. Critical Reviews in Biotechnology 32, 187–202, DOI:10.3109/07388551.2011.595385 (2012).

18 Chundawat, S. P. S., Beckham, G. T., Himmel, M. E. & Dale, B. E. Deconstruction of lignocellulosic biomass to fuels and chemicals. Annu Rev Chem Biomol 2, 121–145, DOI:10.1146/Annurev-Chembioeng-061010-114205 (2011).

19 Himmel, M. E. et al. Biomass recalcitrance: engineering plants and enzymes for biofuels production. Science 315, 804–807, DOI:10.1126/science.1137016 (2007).

20 Payne, C. M. et al. Fungal cellulases. Chem Rev 115, 1308–1448, DOI:10.1021/cr500351c (2015).

21 Lombard, V., Golaconda Ramulu, H., Drula, E., Coutinho, P. M. & Henrissat, B. The carbohydrate-active enzymes database (CAZy) in 2013. Nucleic Acids Research 42, D490–D495, DOI:10.1093/nar/gkt1178 %J Nucleic Acids Research (2013).

22 Sukharnikov, L. O. et al. Sequence, Structure, and Evolution of Cellulases in Glycoside Hydrolase Family 48. 287, 41068–41077, DOI:10.1074/jbc.M112.405720 (2012).

23 Kari, J. et al. A practical approach to steady-state kinetic analysis of cellulases acting on their natural insoluble substrate. Anal Biochem 586, 113411, DOI:10.1016/j.ab.2019.113411 (2019).

24 Kari, J., Andersen, M., Borch, K. & Westh, P. An Inverse Michaelis–Menten Approach for Interfacial Enzyme Kinetics. ACS Catalysis 7, 4904–4914, DOI:10.1021/acscatal.7b00838 (2017).

25 Andersen, M., Kari, J., Borch, K. & Westh, P. Michaelis-Menten equation for degradation of insoluble substrate. Math Biosci 296, 93–97, DOI:10.1016/j.mbs.2017.11.011 (2018).

26 Sousa, S. F., Ramos, M. J., Lim, C. & Fernandes, P. A. Relationship between Enzyme/Substrate Properties and Enzyme Efficiency in Hydrolases. ACS Catalysis 5, 5877–5887, DOI:10.1021/acscatal.5b00923 (2015).

27 Warshel, A. Electrostatic Origin of the Catalytic Power of Enzymes and the Role of Preorganized Active Sites. J. Biol. Chem. 273, 27035–27038, DOI:10.1074/jbc.273.42.27035 (1998).

28 Williams, A. Free Energy Relationships in Organic and Bio-Organic Chemistry. (The Royal Society of Chemistry, 2003).

29 Fersht, A. R., Leatherbarrow, R. J. & Wells, T. N. C. Structure-activity relationships in engineered proteins: analysis of use of binding energy by linear free energy relationships. Biochemistry 26, 6030–6038, DOI:10.1021/bi00393a013 (1987).

30 Fersht, A. R., Leatherbarrow, R. J. & Wells, T. N. C. Quantitative analysis of structure–activity relationships in engineered proteins by linear free-energy relationships. Nature 322, 284–286, DOI:10.1038/322284a0 (1986).

31 Fersht, A. R. & Sato, S. Phi-value analysis and the nature of protein-folding transition states. Proc Natl Acad Sci U S A 101, 7976–7981, DOI:10.1073/pnas.0402684101 (2004).

32 Kurasin, M. & Valjamae, P. Processivity of cellobiohydrolases is limited by the substrate. The Journal of biological chemistry 286, 169–177, DOI:10.1074/jbc.M110.161059 (2011).

33 Cruys-Bagger, N., Tatsumi, H., Ren, G. R., Borch, K. & Westh, P. Transient kinetics and rate-limiting steps for the processive cellobiohydrolase Cel7A: effects of substrate structure and carbohydrate binding domain. Biochemistry 52, 8938–8948, DOI:10.1021/bi401210n (2013).

34 Kipper, K., Väljamäe, P. & Johansson, G. Processive action of cellobiohydrolase Cel7A from Trichoderma reesei is revealed as ‘burst’ kinetics on fluorescent polymeric model substrates. Biochem. J. 385, 527–535, DOI:10.1042/bj20041144 (2005).

35 Murphy, L. et al. Origin of initial burst in activity for Trichoderma reesei endo-glucanases hydrolyzing insoluble cellulose. The Journal of biological chemistry 287, 1252–1260, DOI:10.1074/jbc.M111.276485 (2012).

36 Christensen, S. J., Kari, J., Badino, S. F., Borch, K. & Westh, P. Rate-limiting step and substrate accessibility of cellobiohydrolase Cel6A from Trichoderma reesei. The FEBS journal 285, 4482–4493, DOI:10.1111/febs.14668 (2018).

37 Sørensen, T. H. et al. Selective pressure on an interfacial enzyme: Functional roles of a highly conserved asparagine residue in a cellulase. Biochimica et Biophysica Acta (BBA) – Proteins and Proteomics 1868, 140359, DOI:https://doi.org/10.1016/j.bbapap.2019.140359 (2020).

38 Horn, S. J. et al. Costs and benefits of processivity in enzymatic degradation of recalcitrant polysaccharides. P Natl Acad Sci USA 103, 18089–18094, DOI:10.1073/pnas.0608909103 (2006).

39 Beckham, G. T. et al. Molecular-level origins of biomass recalcitrance: decrystallization free energies for four common cellulose polymorphs. The journal of physical chemistry. B 115, 4118–4127, DOI:10.1021/jp1106394 (2011).

40 Bergenstråhle, M., Thormann, E., Nordgren, N. & Berglund, L. A. Force Pulling of Single Cellulose Chains at the Crystalline Cellulose-Liquid Interface: A Molecular Dynamics Study. Langmuir 25, 4635–4642, DOI:10.1021/la803915c (2009).

41 Payne, C. M. et al. Glycoside hydrolase processivity is directly related to oligosaccharide binding free energy. J Am Chem Soc 135, 18831–18839, DOI:10.1021/ja407287f (2013).

42 Kari, J. et al. Sabatier Principle for Interfacial (Heterogeneous) Enzyme Catalysis. ACS Catalysis 8, 11966–11972, DOI:10.1021/acscatal.8b03547 (2018).

43 Sabatier, P. La catalyse en chimie organique. Vol. III (Librairie Polytechnique, 1913).

44 Bligaard, T. et al. The Brønsted–Evans–Polanyi relation and the volcano curve in heterogeneous catalysis. Journal of Catalysis 224, 206–217 (2004).

45 Nørskov, J. K., Studt, F., Abild-Pedersen, F. & Bligaard, T. Fundamental concepts in heterogeneous catalysis. (John Wiley & Sons, 2014).

46 Borch, K. et al. (Google Patents, 2014).

47 Sorensen, T. H. et al. Temperature Effects on Kinetic Parameters and Substrate Affinity of Cel7A Cellobiohydrolases. The Journal of biological chemistry 290, 22193–22202, DOI:10.1074/jbc.M115.658930 (2015).

48 Pace, C. N., Vajdos, F., Fee, L., Grimsley, G. & Gray, T. How to measure and predict the molar absorption coefficient of a protein. 4, 2411–2423, DOI:10.1002/pro.5560041120 (1995).

49 Trott, O. & Olson, A. J. AutoDock Vina: improving the speed and accuracy of docking with a new scoring function, efficient optimization, and multithreading. Journal of computational chemistry 31, 455–461, DOI:10.1002/jcc.21334 (2010).

50 Guvench, O. et al. CHARMM Additive All-Atom Force Field for Carbohydrate Derivatives and Its Utility in Polysaccharide and Carbohydrate–Protein Modeling. Journal of Chemical Theory and Computation 7, 3162–3180, DOI:10.1021/ct200328p (2011).

51 Abraham, M. J. et al. GROMACS: High performance molecular simulations through multi-level parallelism from laptops to supercomputers. SoftwareX 1-2, 19–25, DOI:https://doi.org/10.1016/j.softx.2015.06.001 (2015).

52 Webb, B. & Sali, A. Comparative Protein Structure Modeling Using MODELLER. 54, 5.6.1-5.6.37, DOI:10.1002/cpbi.3 (2016).

53 Gomes, T. C. F. & Skaf, M. S. Cellulose-Builder: A toolkit for building crystalline structures of cellulose. 33, 1338–1346, DOI:10.1002/jcc.22959 (2012).

54 Beckham, G. T. et al. Identification of amino acids responsible for processivity in a Family 1 carbohydrate-binding module from a fungal cellulase. The journal of physical chemistry. B 114, 1447–1453 (2010).

55 Hub, J. S., de Groot, B. L. & van der Spoel, D. g_wham—A Free Weighted Histogram Analysis Implementation Including Robust Error and Autocorrelation Estimates. Journal of Chemical Theory and Computation 6, 3713–3720, DOI:10.1021/ct100494z (2010).

56 Di Palma, F., Bottaro, S. & Bussi, G. Kissing loop interaction in adenine riboswitch: insights from umbrella sampling simulations. BMC Bioinformatics 16, S6, DOI:10.1186/1471-2105-16-S9-S6 (2015).

57 Akhshi, P. & Wu, G. Umbrella sampling molecular dynamics simulations reveal concerted ion movement through G-quadruplex DNA channels. Physical Chemistry Chemical Physics 19, 11017–11025, DOI:10.1039/C7CP01028A (2017).

58 Patel, J. S. & Ytreberg, F. M. Fast Calculation of Protein–Protein Binding Free Energies Using Umbrella Sampling with a Coarse-Grained Model. Journal of Chemical Theory and Computation 14, 991–997, DOI:10.1021/acs.jctc.7b00660 (2018).

