## Supplementary material for "Physical constrains and functional plasticity of cellulases: Linear scaling relationships for a heterogeneous enzyme reaction": sup. mat.

[1]Department of Biotechnology and Biomedicine, Technical University of Denmark, Søltofts Plads, 2800 Kongens Lyngby, Denmark, [2]Department of Science and Environment, Roskilde University, Universitetsvej 1, 4000 Roskilde, Denmark, [3]Novozymes A/S, Krogshøjvej 36, 2880 Bagsværd, Denmark, [4]Department of Geosciences and Natural Resource Management, University of Copenhagen, Rolighedsvej 23, 1958 Frederiksberg C, Denmark, [5]Department of Chemistry, Technical University of Denmark, Kemitorvet 207, 2800 Kongens Lyngby, Denmark

*[#] Both authors contributed equally to this work*

### Supplementary Methods

##### 1. Derivation of Optimal Michealis-Menten Constant ( $K_M^{opt}$ )

Equation 6 in the main text (Eq. S1) shows how $K_M$ determines the initial rate.

$$v = \frac{E_0 A K_m{}^a S_0}{S_0 + K_M} \qquad \text{(Eq. S1)}$$

The optimal $K_M$ can be found by solving the equation $\frac{dV}{dK_M} = 0$ for $K_M$. Using the quotient rule $\frac{d}{dx}\left(\frac{u}{v}\right) = \frac{v\frac{du}{dx} - u\frac{dv}{dx}}{v^2}$ the derivative of Eq. S1 can be calculated

$$\frac{dV / E_0}{dK_M} = \frac{S_0 A K_m{}^{a-1}(a(S_0 + K_M) - K_M)}{(S_0 + K_M)^2} = 0$$

Solving for $K_M$ gives

$$K_M^{opt} = S_0 \frac{a}{1-a} \qquad \text{(Eq. S2)}$$

Using Eq. S1 and S2 we simulated volcano-curves and their Sabatier optimum for 50 substrate loads ranging from 1 to 100 g/L (Supplementary Figure 1).

#### 2. Insoluble reducing ends measurements

All calculated rates in the main manuscript are based on the amount of soluble product produced. Hence, the endolytic activity of EGs is not accounted for in the analysis and this could lead to an underestimation of the EGs activities if they have a significant production of insoluble products (e.g. insoluble cellooligosaccharides or new chain ends). In order to quantify possible systematic underestimations of the activities of the EGs, we measured the amount of insoluble reducing ends for a representative group of enzymes. The group included EGs from all of the investigated GH families and a CBH enzyme from family 6 and 7. In general, the results showed that that the insoluble activity was small compared to the soluble activity (see table S1). Only TrCel12A had a significant insoluble activity (46%), which may explain why this enzyme appeared at the edge of the 95% prediction band in Fig. 2 (main text).
The results in table S1 were estimated by adapting the method described in Silveira, et al. [1] and by using Bicinchoninic acid (BCA) in presence of $CuSO_4$. The reactions were carried out in 2 mL microcentrifuge tubes (Sarstedt, Numbrecht, Germany). Enzymes and washed Avicel (Avicel PH-101, Sigma-Aldrich, Steinheim, Germany) were mixed in 300 µL to a final enzyme concentration of 0.1 µM and Avicel load of 40 g/L. The reactions were incubated for 1h at 25 °C in thermomixers (Eppendorf, Hamburg, Germany) equipped with ThermoTop and operating at 1100 rpm. The reactions were then stopped by transfer on ice where the rest of the procedure followed. From the reactions, 25 µL were retrieved and standard buffer was added to a final volume of 500 µL. These samples were used to quantify the total reducing ends (TRE). The initial reactions were then centrifuged for 3 min at 14100 rcf. From these, 25

μL were retrieved and standard buffer was added to a final volume of 500 μL. The samples were centrifuged again as described before. These samples were used to measure the amount of soluble reducing ends (SRE). The amount of insoluble reducing ends where calculated by the subtracting SRE from the TRE. A volume of 150 μL was retrieved from both TRE and SRE samples, and 150 μL of Bicinchoninic acid (BCA) reagent was added. BCA reagent was prepared by mixing equal volumes of solution A (0.46 M sodium carbonate, 0.26 M sodium bicarbonate and 4.5 mM Bicinchoninic acid disodium salt hydrate in deionized water) and solution B ( 4.5 mM Copper(II) sulfate pentahydrate and 10 mM L-serine in deionized water). The samples were briefly vortexed and incubated in thermomixers (Eppendorf, Hamburg, Germany) for 30 min at 75 °C 1100 rpm. The reactions were then cooled for 10 min at 4 °C, vortexed again, and centrifuged for 3 min as described above. Finally, 100 μL of supernatant were transferred to a 96 well microtiter plate (655101, Greiner Bio-One, Germany) and the absorbance at 560 nm was measured using a spectrophotometer (Spectramax i3, Molceular Devices, Wals, Austria). A 6-point calibration curve of cellobiose (125-3.9 μM) was included in all measurements. Results from the above analysis is reported in table S1 for a selected group of cellulases. The group was selected so that all types of endoglucanases and cellobiohydrolases was covered.

### Supplementary tables

**Supplementary Table 1.** Concentration of total reducing end (TRE), soluble reducing ends (SRE) and insoluble reducing ends (IRE) for 4 different endoglucanases (EG) and 2 different cellobiohydrolases (CBH)

| Enzyme | Mode of action | TRE (µM) | SRE (µM) | IRE (µM) | IRE/TRE *100 |
|---|---|---|---|---|---|
| TrCel5A | EG | 169 ±30 | 156 ±3 | 13 ±30 | 8 |
| TrCel12A | EG | 145 ±56 | 78 ±1.5 | 67 ±56 | 46 |
| HiCel45A | EG | 131 ±9 | 124.5 ±0.4 | 6 ±9 | 5 |
| TrCel7B | EG | 220 ±28 | 211 ±2 | 9 ±28 | 4 |
| TrCel7A | CBH | 35 ±15 | 36 ±6 | n.d. | n.d. |
| TrCel6A | CBH | 98 ±63 | 103 ±4 | n.d. | n.d. |

**Supplementary Table 2.** Wild-type and variant enzymes properties. "O", catalytic domain; "o", CBM1; "---", linker. Blue indicates mutations in the catalytic domain, purple in the CBM, and yellow in the linker. **Bold** indicates chimeric enzymes where a CBM were added or substituted.

| Organism | GH | Mechanism | WT or Variant | Modularity | Mutations | Sequence information[1] | Ref.[2] | $k_{cat}$ ($s^{-1}$) | $K_M$ (g/L) | $(k_{cat}/K_M)*10^3$ ($s^{-1}gL^{-1}$) |
|---|---|---|---|---|---|---|---|---|---|---|
| **Cellobiohydrolases** | | | | | | | | | | |
| *Trichoderma reesei* | 7 | Retaining | WT | O---o | N/A | P62694 (U) | 2 | 0.095 ± 0.003 | 2.6 ± 0.3 | 35.8 ±2.6 |
| *Penicillium chrysogenum* | 7 | Retaining | WT | O---o | N/A | Q5S1P9 (U) | | 0.388 ± 0.006 | 22.4 ± 1.0 | 17.3 ±0.5 |
| *Talaromyces leycettanus* | 7 | Retaining | WT | O---o | N/A | S6EXC0 (U) | | 0.076 ± 0.003 | 1.2 ± 0.1 | 62.6 ±4.9 |
| *Rasamsonia emersonii* | 7 | Retaining | WT | O | N/A | Q8TFL9 (U) | 3 | 0.091 ± 0.003 | 6.0 ± 0.7 | 15 ±1.1 |
| *Rasamsonia byssoclamydoides* | 7 | Retaining | WT | O | N/A | S6EJQ7 (U) | | 0.112 ± 0.008 | 7.6 ± 1.3 | 14.7 ±1.5 |
| *Neosartorya fischeri* | 7 | Retaining | WT | O | N/A | A1DMA5 (U) | | 1.354 ± 0.119 | 115.1 ± 15.7 | 11.8 ±0.6 |
| *Neosartorya fischeri* | 7 | Retaining | WT | O---o | N/A | A1DAP8 (U) | | 0.106 ± 0.003 | 1.6 ± 0.2 | 65.4 ±4.7 |
| *Aspergillus terreus* | 7 | Retaining | WT | O---o | N/A | Q0CMT2 (U) | | 0.216 ± 0.006 | 6.4 ± 0.6 | 33.9 ±2.5 |
| *Colletotrichum graminicola* | 7 | Retaining | WT | O | N/A | E3Q986 (U) | | 0.094 ± 0.003 | 4.8 ± 0.5 | 19.6 ±1.6 |
| *Phanerochaete chrysosporium* | 7 | Retaining | WT | O---o | N/A | P13860 (U) | | 0.143 ± 0.004 | 3.5 ± 0.3 | 40.6 ±2.4 |
| *Aspergillus aculeatus* | 7 | Retaining | WT | O | N/A | A0A1L9X3D1 (U) | | *0.251 ± 0.014 | *41.5 ± 5.2 | *6.0 ±0.4 |
| *Trichoderma reesei* | 6 | Inverting | WT | o---O | N/A | P07987 (U) | 4 | 0.564 ± 0.034 | 26.4 ± 4.0 | 21.4 ±1.9 |
| *Lentinus sajor-caju* | 6 | Inverting | WT | o---O | N/A | Q96TP4 (U) | 4 | 1.345 ± 0.060 | 101.0 ± 7.5 | 13.3 ±0.4 |
| *Aspergillus terreus* | 6 | Inverting | WT | O | N/A | Q0D1J1 (U) | 4 | 4.900 ± 0.785 | 708.3 ± 127.1 | 6.9 ±0.1 |
| *Colletotrichum graminicola* | 6 | Inverting | WT | o---O | N/A | E3Q540 (U) | 4 | 1.172 ± 0.178 | 65.2 ± 19.0 | 18 ±2.5 |
| *Colletotrichum graminicola* | 6 | Inverting | WT | O | N/A | E3Q986 (U) | 4 | U/D | U/D | 9.4 ±0.1 |
| *Trichoderma reesei* | 7 | Retaining | Variant | O | Deletion of linker and CBM | N/A | 2 | 0.165 ± 0.003 | 8.7 ± 0.6 | 19.1 ±0.9 |
| *Trichoderma reesei* | 7 | Retaining | Variant | O---o---**o** | CBM and linker from TrCel7A | N/A | | 0.087 ± 0.002 | 2.5 ± 0.2 | 34.4 ±2 |
| *Trichoderma reesei* | 7 | Retaining | Variant | O---o | W38A | N/A | 2 | 0.506 ± 0.056 | 35.1 ± 9.0 | 14.4 ±2.1 |
| *Trichoderma reesei* | 7 | Retaining | Variant | O---o | W40A | N/A | 5 | 0.134 ± 0.003 | 4.2 ± 0.4 | 31.9 ±2.3 |
| *Trichoderma reesei* | 7 | Retaining | Variant | O---o | T246C/Y371C | N/A | 6 | 0.094 ± 0.003 | 5.4 ± 0.6 | 17.3 ±1.2 |
| *Trichoderma reesei* | 7 | Retaining | Variant | O---o | Δ(W192-G205) | N/A | 7 | 0.347 ± 0.025 | 16.0 ± 3.3 | 21.7 ±3 |
| *Trichoderma reesei* | 7 | Retaining | Variant | O---o | Δ(E193-G205) | N/A | 7 | 0.363 ± 0.022 | 16.4 ± 2.9 | 22.1 ±2.6 |
| *Trichoderma reesei* | 7 | Retaining | Variant | O---o | Δ(S196-T201) | N/A | 7 | 0.287 ± 0.007 | 11.7 ± 0.9 | 24.5 ±1.3 |
| *Trichoderma reesei* | 7 | Retaining | Variant | O-o | Δ(G439-G444) | N/A | | 0.071 ± 0.002 | 1.6 ± 0.2 | 45.9 ±3.3 |
| *Rasamsonia emersonii* | 7 | Retaining | Variant | O---**o** | CBM and linker from *T. reesei* Cel7A | N/A | 3 | 0.060 ± 0.001 | 1.0 ± 0.1 | 60 ±5 |
| *Rasamsonia emersonii* | 7 | Retaining | Variant | O---**o** | CBM and linker from *T. reesei* Cel7A + Y470W | N/A | | 0.050 ± 0.003 | 1.0 ± 0.2 | 48.4 ±6.2 |
| *Rasamsonia emersonii* | 7 | Retaining | Variant | O---**o** | CBM and linker from *T. reesei* Cel7A + Y478W | N/A | | 0.093 ± 0.002 | 5.0 ± 0.4 | 18.7 ±1.2 |
| *Rasamsonia emersonii* | 7 | Retaining | Variant | O---**o** | CBM and linker from *T. reesei* Cel7A + Y478W/Y497W | N/A | | 0.077 ± 0.001 | 3.7 ± 0.2 | 20.9 ±1 |
| *Rasamsonia emersonii* | 7 | Retaining | Variant | O---**o** | CBM and linker from *T. reesei* Cel7A + Y496W/Y497W | N/A | | 0.063 ± 0.003 | 2.0 ± 0.3 | 31.7 ±2.7 |
| *Rasamsonia emersonii* | 7 | Retaining | Variant | O---**o** | CBM and linker from *T. reesei* Cel7A + Y470W/Y478W | N/A | | 0.088 ± 0.002 | 3.3 ± 0.2 | 26.6 ±1.4 |
| *Rasamsonia emersonii* | 7 | Retaining | Variant | O---**o** | CBM and linker from *T. reesei* Cel7A + Y478W/Y496W | N/A | | 0.092 ± 0.003 | 4.9 ± 0.6 | 18.7 ±1.6 |
| *Rasamsonia emersonii* | 7 | Retaining | Variant | O---**o** | CBM and linker from *T. reesei* Cel7A + Y470W/Y496W | N/A | | 0.086 ± 0.002 | 4.9 ± 0.3 | 17.6 ±0.7 |
| *Rasamsonia emersonii* | 7 | Retaining | Variant | O---**o** | CBM and linker from *T. reesei* Cel7A + Y470W/Y496W/Y497W | N/A | | 0.083 ± 0.002 | 2.9 ± 0.2 | 28.8 ±1.6 |
| *Rasamsonia emersonii* | 7 | Retaining | Variant | O---**o** | CBM and linker from *T. reesei* Cel7A + Y478W/Y496W/Y497W | N/A | | 0.087 ± 0.002 | 3.1 ± 0.3 | 27.8 ±1.6 |
| *Rasamsonia emersonii* | 7 | Retaining | Variant | O---**o** | CBM and linker from *T. reesei* Cel7A + Y470W/Y478W/Y496W/Y497W | N/A | | 0.075 ± 0.004 | 3.8 ± 0.7 | 19.7 ±2.3 |
| *Rasamsonia emersonii* | 7 | Retaining | Variant | O---**o** | CBM and linker from *T. reesei* Cel7A + Y470W/Y478W/Y496W | N/A | | 0.078 ± 0.002 | 4.4 ± 0.4 | 18 ±1.2 |
| *Trichoderma reesei* | 6 | Inverting | Variant | O | Deletion of linker and CBM | N/A | 8 | 0.834 ± 0.047 | 58.7 ± 6.5 | 14.2 ±0.8 |
| *Trichoderma reesei* | 6 | Inverting | Variant | o---O | Y103A | N/A | | 0.618 ± 0.054 | 30.5 ± 6.4 | 20.3 ±2.5 |
| *Trichoderma reesei* | 6 | Inverting | Variant | o---O | N305A | N/A | | 0.856 ± 0.071 | 44.0 ± 7.7 | 19.5 ±1.8 |
| *Trichoderma reesei* | 6 | Inverting | Variant | o---O | S106A | N/A | | 0.697 ± 0.064 | 36.0 ± 7.6 | 19.4 ±2.3 |
| *Trichoderma reesei* | 6 | Inverting | Variant | o---O | R410A | N/A | | 0.524 ± 0.044 | 30.3 ±6.1 | 17.3 ±2 |
| *Trichoderma reesei* | 6 | Inverting | Variant | o---O | G365D/D366N | N/A | | 0.637 ± 0.069 | 43.1 ± 10.0 | 14.8 ±1.8 |
| *Trichoderma reesei* | 6 | Inverting | Variant | o---O | W367F | N/A | | 1.087 ± 0.204 | 66.6 ± 23.8 | 16.3 ±2.8 |
| *Trichoderma reesei* | 6 | Inverting | Variant | o---O | W269A | N/A | | 1.578 ± 0.216 | 156.4 ± 30.7 | 10.1 ±0.6 |
| *Trichoderma reesei* | 6 | Inverting | Variant | o---O | W272A | N/A | | 1.809 ± 0.268 | 110.0 ± 26.5 | 16.4 ±1.5 |
| *Trichoderma reesei* | 6 | Inverting | Variant | o---O | K395T | N/A | | *0.015 ± 0.003 | *15.3 ± 8.9 | *1.0 ±0.4 |
| *Trichoderma reesei* | 6 | Inverting | Variant | o---O | W269A/W272A | N/A | | *0.629 ± 0.198 | *159.0 ± 74.1 | *4.0 ±0.6 |
| *Trichoderma reesei* | 6 | Inverting | Variant | **o**---O | CBM from *P. anserina* GH6 | N/A | 9 | 0.514 ± 0.028 | 28.8 ± 3.8 | 17.8 ±1.4 |
| *Trichoderma reesei* | 6 | Inverting | Variant | **o**---O | CBM from *N. frontalis* GH6 | N/A | 9 | 0.564 ± 0.028 | 29.6 ± 3.5 | 19.1 ±1.3 |
| *Trichoderma reesei* | 6 | Inverting | Variant | **o**---O | CBM from *S. indica* GH6 | N/A | 9 | 0.391 ± 0.020 | 28.2 ± 3.6 | 13.9 ±1.0 |
| *Trichoderma reesei* | 6 | Inverting | Variant | **o**---O | CBM from *C. cinerea* GH6 | N/A | 9 | 0.770 ± 0.042 | 45.1 ± 5.1 | 17.1 ±1.0 |
| *Trichoderma reesei* | 6 | Inverting | Variant | **o**---O | CBM from *V. volvacea* GH6 | N/A | 9 | 0.654 ± 0.040 | 32.4 ± 4.7 | 20.2 ±1.7 |
| **Endoglucanases** | | | | | | | | | | |
| *Trichoderma reesei* | 7 | Retaining | WT | O---o | N/A | P07981 (U) | 7 | 2.365 ± 0.243 | 115.2 ± 18.4 | 20.5 ±1.2 |
| *Chaetomium virescens* | 7 | Retaining | WT | O | N/A | BDB18317 (GS) | | 5.360 ± 0.800 | 782.9± 128.1 | 6.8 ±0.1 |
| *Aspergillus terreus* | 7 | Retaining | WT | O | N/A | Q0CC84 (U) | | U/D | U/D | 5.0 ±0.1 |
| *Trichoderma reesei* | 5 | Retaining | WT | O---o | N/A | P07982 (U) | 10 | 1.150 ± 0.077 | 58.9 ± 7.6 | 19.5 ±1.2 |
| *Thermoascus aurantiacus* | 5 | Retaining | WT | O | N/A | Q8TG26 (U) | | 2.500 ± 0.553 | 352.9 ± 94.1 | 7.1 ±0.3 |
| *Penicillium brasilianum* | 5 | Retaining | WT | O---o | N/A | B8Q961 (U) | | 0.785 ± 0.038 | 45.7 ± 4.6 | 17.2 ±0.9 |
| *Neosartorya fischeri* | 5 | Retaining | WT | O---o | N/A | A1DNK9 (U) | | 0.935 ± 0.070 | 62.5 ± 8.8 | 15 ±1 |
| *Gloephyllum trabeum* | 5 | Retaining | WT | O | N/A | D7REW1 (U) | | 0.705 ± 0.061 | 53.8 ± 9.2 | 13.1 ±1.1 |
| *Chaetomium virescens* | 5 | Retaining | WT | O | N/A | BDB1821(GS) | | 1.086 ± 0.119 | 91.4 ± 16.7 | 11.9 ±0.9 |
| *Colletotrichum graminicola* | 5 | Retaining | WT | o---O | N/A | E3QRW7 (U) | | 0.938 ± 0.057 | 84.2 ± 8.7 | 11.1 ±0.5 |
| *Aspergillus aculeatus* | 5 | Retaining | WT | O | N/A | A0A1L9WYV6 (U) | | Activity too low to be measured | | |
| *Humicola insolens* | 45 | Inverting | WT | O---o | N/A | P43316 (U) | | 1.265 ± 0.102 | 97.0 ± 12.9 | 13 ±0.7 |
| *Humicola hyalothermophila* | 45 | Inverting | WT | O---o | N/A | BBJ30934 (GS) | | 0.884 ± 0.027 | 80.8 ± 4.3 | 10.9 ±0.2 |
| *Acremonium furcatum* | 45 | Inverting | WT | O---o | N/A | AAY00847.1 (GB) | | 0.664 ± 0.016 | 42.3 ± 2.1 | 15.7 ±0.4 |
| *Lectera colletotrichoides* | 45 | Inverting | WT | O---o | N/A | AAY00854.1 (GB) | | 0.495 ± 0.010 | 33.8 ± 1.6 | 14.6 ±0.4 |
| *Rhizomucor pusillus* | 45 | Inverting | WT | O---o | N/A | BBK79186 (GS) | | 0.571 ± 0.024 | 69.2 ± 5.4 | 8.3 ±0.3 |
| *Melanocarpus albomyces* | 45 | Inverting | WT | O | N/A | Q8J0K8 (U) | | *0.152 ± 0.013 | *19.4 ± 4.7 | *7.8 ±1.2 |
| *Trichoderma reesei* | 12 | Retaining | WT | O | N/A | G0RRG8 (U) | 11 | 0.718 ± 0.044 | 102.9 ± 10.2 | 7.0 ±0.3 |
| *Aspergillus aculeatus* | 12 | Retaining | WT | O | N/A | A0A1L9WFV2 (U) | | Activity too low to be measured | | |
| *Chaetomium virescens* | 12 | Retaining | WT | O | N/A | BDB18192 (GS) | | *0.147 ± 0.041 | *86.0 ± 40.8 | *1.7 ±0.3 |
| *Trichoderma reesei* | 7 | Retaining | Variant | O---o | D76R | N/A | | 0.947 ± 0.052 | 59.7 ± 6.2 | 15.9 ±0.8 |
| *Trichoderma reesei* | 7 | Retaining | Variant | O---o | D62A/E63A | N/A | | 0.308 ± 0.010 | 31.1 ± 2.4 | 9.9 ±0.4 |
| *Trichoderma reesei* | 7 | Retaining | Variant | O---o | D115A | N/A | | 1.502 ± 0.169 | 84.0 ± 16.7 | 17.9 ±1.5 |
| *Trichoderma reesei* | 7 | Retaining | Variant | O---o | D151A, E152A | N/A | | 1.135 ± 0.084 | 77.3 ± 10.4 | 14.7 ±0.9 |
| *Trichoderma reesei* | 7 | Retaining | Variant | O---o | D366P | N/A | | 1.404 ± 0.120 | 80.6 ± 11.9 | 17.4 ±1.1 |
| *Trichoderma reesei* | 7 | Retaining | Variant | O---o | K122T | N/A | | 1.564 ± 0.130 | 99.5 ± 13.4 | 15.7 ±0.8 |
| *Trichoderma reesei* | 12 | Retaining | Variant | O | Y111W | N/A | | 0.792 ± 0.090 | 115.4 ± 21.0 | 6.9 ±0.5 |
| *Trichoderma reesei* | 12 | Retaining | Variant | O | S63I | N/A | | 0.783 ± 0.076 | 101.9 ± 16.4 | 7.7 ±0.5 |
| *Trichoderma reesei* | 12 | Retaining | Variant | O | W7G | N/A | | *0.445 ± 0.134 | *238.0 ± 95.5 | *1.9 ±0.2 |
| *Trichoderma reesei* | 12 | Retaining | Variant | O | W22A | N/A | | Activity too low to be measured | | |

[1]UniProt (U), Genbank (GB) or GENESEQ (GS) accession number.
[2]Enzyme was constructed, expressed and purified as described in the reference.
*Identified as outliers in Fig. 2 in the main text.
**N/A** = not applicable, **U/D** = unable to determine. The affinity of the enzyme was too low to resolve $K_M$ and $k_{cat}$ from non-linear regression to Eq. 1 (main text). Only the ratio $k_{cat}/K_M$ could be determined.

### Supplementary figures

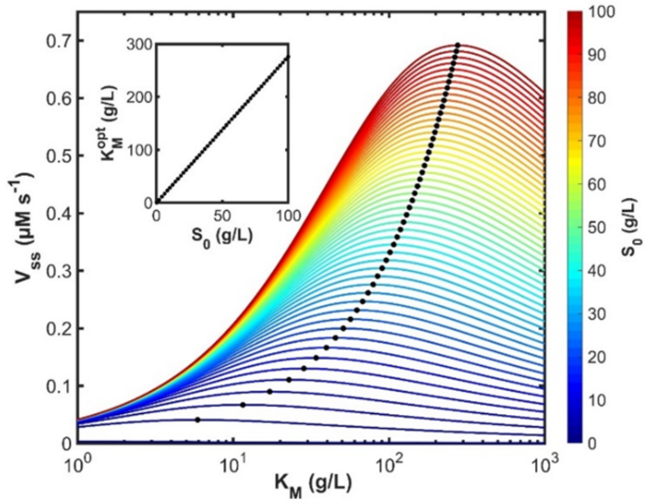

**Supplementary Figure 1**. Simulated volcano curves for 50 different substrate loads ranging from 1 g/L to 100 g/L. The black circles indicate the optimal affinity ( $K_M^{opt}$ ) and the insert shows how $K_M^{opt}$ scales with the substrate load ($S_0$). The curves were simulated with Eq. S1 and $K_M^{opt}$ was predicted using Eq. S2.

### References

1 Silveira, M. H. L., Aguiar, R. S., Siika-aho, M. & Ramos, L. P. Assessment of the enzymatic hydrolysis profile of cellulosic substrates based on reducing sugar release. *Bioresource Technology* **151**, 392-396, doi:https://doi.org/10.1016/j.biortech.2013.09.135 (2014).

2 Kari, J. *et al.* Kinetics of Cellobiohydrolase (Cel7A) Variants with Lowered Substrate Affinity. *The Journal of biological chemistry* **289**, 32459-32468, doi:10.1074/jbc.M114.604264 (2014).

3 Sorensen, T. H. *et al.* Temperature Effects on Kinetic Parameters and Substrate Affinity of Cel7A Cellobiohydrolases. *The Journal of biological chemistry* **290**, 22193-22202, doi:10.1074/jbc.M115.658930 (2015).

4 Christensen, S. J., Krogh, K. B. R. M., Spodsberg, N., Borch, K. & Westh, P. A biochemical comparison of fungal GH6 cellobiohydrolases. *Biochem J* **476**, 2157-2172, doi:10.1042/BCJ20190185 %J Biochemical Journal (2019).

5 Røjel, N. *et al.* Substrate binding in the processive cellulase Cel7A: Transition state of complexation and roles of conserved tryptophan residues. doi:10.1074/jbc.RA119.011420 (2019).

6 Kari, J. *et al.* Sabatier Principle for Interfacial (Heterogeneous) Enzyme Catalysis. *ACS Catalysis* **8**, 11966-11972, doi:10.1021/acscatal.8b03547 (2018).

7 Schiano-di-Cola, C. *et al.* Systematic deletions in the cellobiohydrolase (CBH) Cel7A from the fungus Trichoderma reesei reveal flexible loops critical for CBH activity. *The Journal of biological chemistry* **294**, 1807-1815, doi:10.1074/jbc.RA118.006699 (2019).

8 Badino, S. F. *et al.* Exo-exo synergy between Cel6A and Cel7A from Hypocrea jecorina: Role of carbohydrate binding module and the endo-lytic character of the enzymes. **114**, 1639-1647, doi:10.1002/bit.26276 (2017).

9 Christensen, S. J., Badino, S. F., Cavaleiro, A. M., Borch, K. & Westh, P. Functional analysis of chimeric TrCel6A enzymes with different carbohydrate binding modules. *Protein Engineering, Design and Selection*, doi:10.1093/protein/gzaa003 (2020).

10 Murphy, L. *et al.* Origin of initial burst in activity for Trichoderma reesei endo-glucanases hydrolyzing insoluble cellulose. *The Journal of biological chemistry* **287**, 1252-1260, doi:10.1074/jbc.M111.276485 (2012).

11 Kari, J., Andersen, M., Borch, K. & Westh, P. An Inverse Michaelis–Menten Approach for Interfacial Enzyme Kinetics. *ACS Catalysis* **7**, 4904-4914, doi:10.1021/acscatal.7b00838 (2017).
